# Differential impact of TNFRSF co-stimulation on CD8^+^ T cell cytokine production by feedback control of surface receptor expression

**DOI:** 10.1101/2021.07.02.450833

**Authors:** John Nguyen, Johannes Pettmann, Philipp Kruger, Omer Dushek

## Abstract

T cell responses towards infections and cancers are regulated by a host of co-signalling receptors that are largely grouped into the binary categories of co-stimulation and co-inhibition. The TNF receptor superfamily (TNFRSF) members 4-1BB, CD27, GITR, and OX40 are well-established co-stimulation receptors with largely shared molecular pathways raising the question of whether they also have a similar impact on quantitative T cell responses, such as the efficacy, sensitivity, and duration of T cell responses. Here, we systematically stimulated primary human CD8^+^ T cell blasts with dose ranges of antigen and ligands for TNFRSF members to screen for their quantitative effects on cytokine production. Although both 4-1BB and CD27 increased efficacy, only 4-1BB was able to prolong the duration of cytokine production, and both had only a modest impact on antigen sensitivity. An operational model could explain these divergent quantitative phenotypes using a shared signalling mechanism based on the surface expression of 4-1BB, but not CD27, being regulated through a signalling feedback. The model predicted that CD27 co-stimulation would increase 4-1BB expression and subsequent 4-1BB co-stimulation, which we confirmed experimentally. Although GITR and OX40 produced only minor changes in cytokine production on their own, we found that like 4-1BB, CD27 could enhance GITR expression and subsequent GITR co-stimulation. Thus, feedback control of induced TNFRSF surface expression explains both synergy and differential impact on cytokine production. The work highlights that different co-stimulation receptors can have different quantitative phenotypes on the same output allowing for highly regulated control of T cell responses.

## Introduction

T cells are critical mediators of adaptive immunity against pathogens and tumours. Although their activity is primarily controlled by the T cell receptor (TCR), a range of other co-signalling receptors are also known to significantly modulate the TCR signal and hence T cell activity (1). Depending on their overall positive or negative impact on T cell activity, these co-signalling receptors have been binarily divided into co-stimulatory (e.g. CD28 and 4-1BB) or co-inhibitory receptors (such as CTLA-4 and PD-1). Many of these receptors fall into the immunoglobulin and tumour necrosis factor receptor superfamilies (IgSF and TNFRSF, respectively) which differ in structure and signalling mechanisms. Even within these families, the expression patterns and functional phenotypes of these receptors display a large variety. Despite these differences, our classification of these co-signalling receptors has largely been confined to the binary qualitative categories based on whether ligation of a co-signalling receptor increases (co-stimulatory) or decreases (co-inhibitory) T cell responses.

Critically important quantitative features of a T cell function (such as cytokine production and target killing) include rate, sensitivity, and duration of the response (Fig. 1). For example, increases in the rate of cytokine production in response to a certain amount of presented antigen have been demonstrated for the archetypal co-stimulatory receptor CD28 (2). Enhanced sensitivity, on the other hand, would allow T cells to respond to lower doses of antigen, which is well-established for adhesion receptors (e.g. CD2 and LFA-1) (3–5), although this also appears to be the case for CD28 to some extent (6, 7). Finally, it has been shown in multiple *in vitro* and *in vivo* systems that T cells exhibit adaptation so that they produce cytokine for limited duration to a constant dose of antigen (8–10), and this duration can be controlled by co-signalling receptors (11). Existing evidence suggests that co-stimulatory TNFRSF members may increase the rate of the response (12–14), but their impact on sensitivity and duration of the T cell response is poorly characterised. Quantitative differences in T cell responses may provide a rational for why similar surface receptors are expressed on the same T cell population.

**Figure 1:**
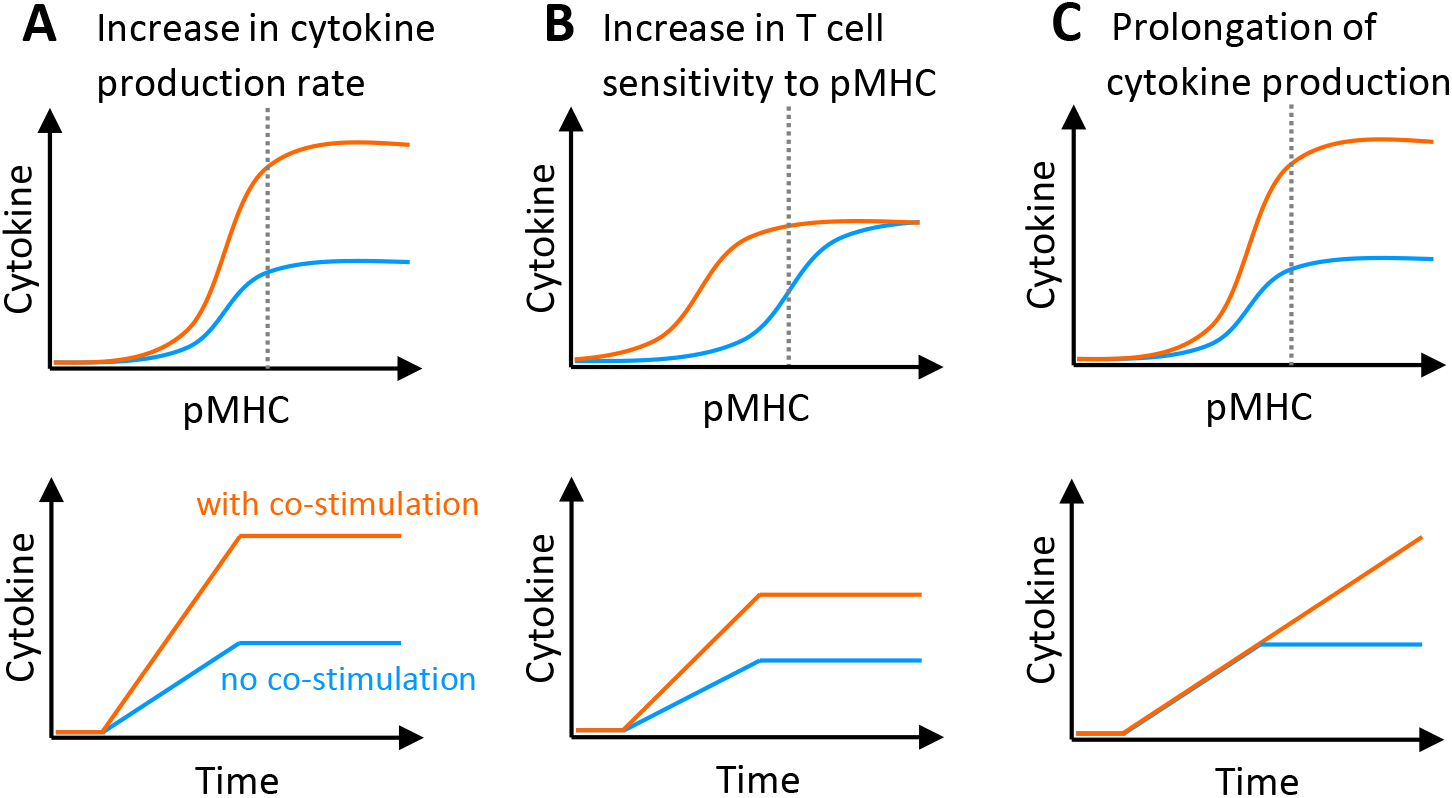
Quantitative effects of co-stimulation on the T cell cytokine response. Graphical representation of hypothetical co-stimulation affecting the (A) rate, (B) sensitivity and (C) duration of the T cell cytokine response, shown as pMHC dose-response at the endpoint (top row) and time courses at the pMHC dose indicated by the dotted line (bottom row). The y-axis represents the cumulative amount of cytokine produced. Blue and orange lines represent the cytokine response in absence and presence of co-stimulation, respectively.

Here we focus on TNF receptor-associated factor (TRAF)-binding receptors of the TNFRSF, with 4-1BB (CD137), CD27, OX40 (CD134) and GITR (AITR, CD357) as well-established representative co-stimulatory receptors on T cells. Their signalling mechanisms and components of their molecular pathways have been characterised in great detail (15, 16). At the same time, a range of functional data has been collected for these receptors in various *in vitro* and *in vivo* systems and in the clinic (17), often focusing on qualitative features of co-stimulation such as enhanced T cell proliferation, survival, anti-viral or anti-tumour response, differentiation, and memory formation. This wealth of information is now being utilised for the development of biotherapeutics targeting TNFRSF members (18), as well as adoptive cell therapies for the treatment of cancers using T cells engineered with mechanisms to activate their pathways (19). Despite previous breakthroughs (20), these promising endeavours are often impeded by modest performance in clinical trials (21–23). Although our molecular understanding of these receptors is mature, our “operational” understanding (24) of how they quantitatively shape T cell responses, including their ability to control the rate and duration of cytokine production, and their impact on antigen sensitivity, is underexplored. This information can improve our operational understanding of these receptors, which can in turn improve our ability to predict T cell responses when these receptors are targeted in therapy.

Using primary human CD8^+^ T cells, we systematically explored the impact of TNFRSF co-stimulation on quantitative T cell responses. We found that 4-1BB and CD27 increased the rate of cytokine production but only 4-1BB could also prolong the duration, and both receptors had only a modest impact on antigen sensitivity. Ligands to GITR and OX40 had only modest effects, possibly due to their relatively low expression on CD8^+^ T cells. The systematic quantitative data allowed us to construct a mathematical model that reconciled these different phenotypes with their largely shared signalling mechanisms by relying on differences in the regulation of surface receptor expression. This operational model predicted a synergy between the receptors based on feedback control of surface expression of inducible TNFRSF members and we confirmed this to be the case by showing that CD27 co-stimulation improved subsequent co-stimulation not only by 4-1BB, but also by GITR. The work highlights how T cell co-stimulation even by similar surface receptors can exhibit differences in quantitative responses and synergy.

## Results

### Co-stimulation through TNFRSF members produces quantitatively different phenotypes

To quantitatively study TNFRSF co-stimulation, we first isolated, *in vitro* expanded, and transduced primary human CD8^+^ T cells with the c58c61 TCR (25) using a standard adoptive T cell therapy protocol (26). This affinity-enhanced TCR recognises the cancer-testes antigen NY-ESO-1_157-165_ on HLA-A2, of which we used a variant with physiological affinity (K_D_ = 7.2 µM (27), see also Materials & Methods). To precisely control pMHC antigen and TNFRSF ligand dose and duration of stimulation (28), these T cells were presented with recombinant ligands on plates (27, 29–31) (Fig. 2A). We systematically stimulated T cells with 12 doses of pMHC and 3-7 doses of trimeric ligands to four members of the TNFRSF; 4-1BB, CD27, GITR and OX40, to study the impact of TNFRSF co-stimulation on quantitative T cell responses. By measuring T cell responses at 4 different time points, we generated a dataset with 1056 independent conditions (Fig. 2B). This allowed us to accurately determine the maximal efficacy of cytokine production (E_max_, maximal response across different antigen doses) and antigen sensitivity (EC_50_, antigen dose at which half-maximal response is observed) at different time points.

**Figure 2:**
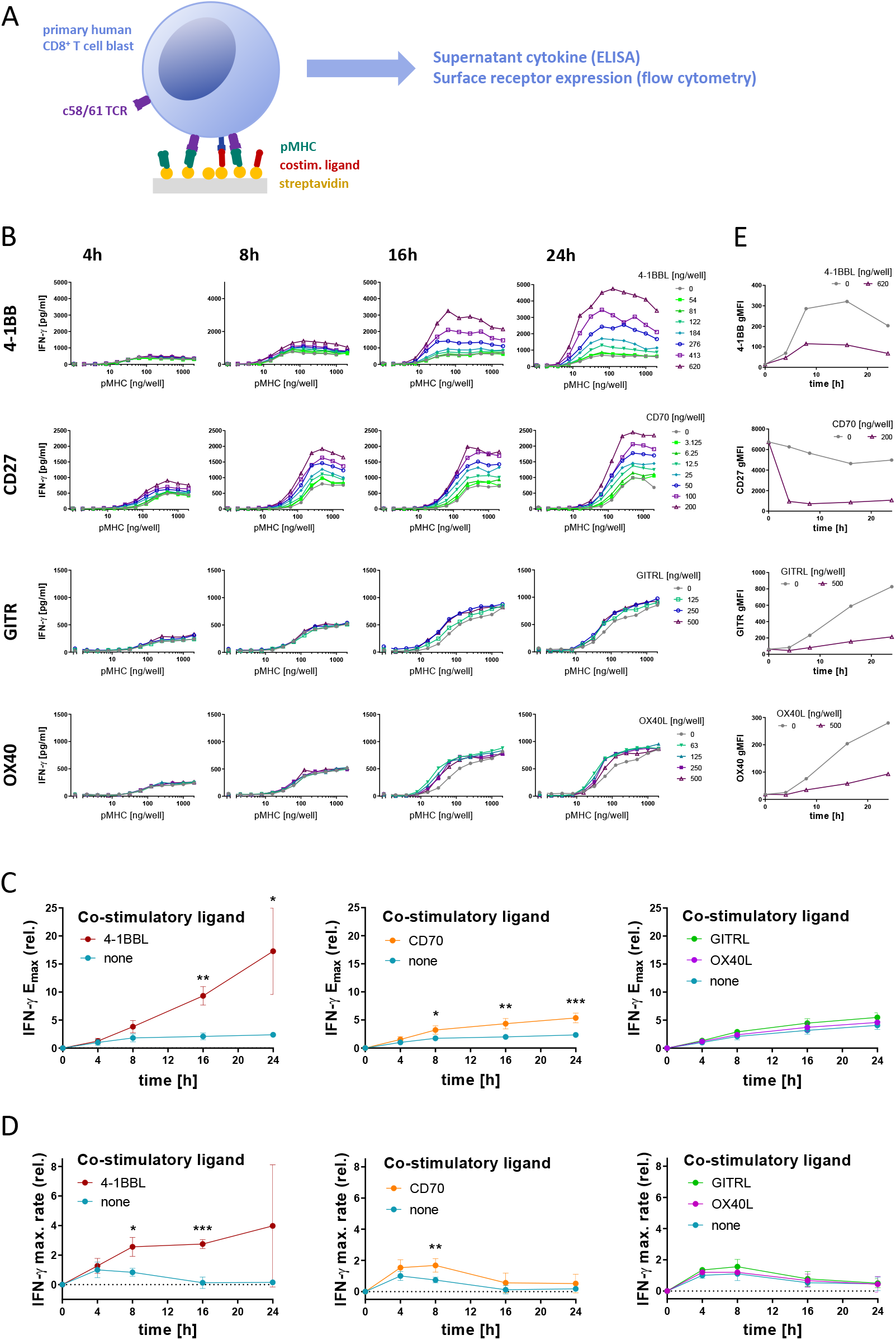
CD8^+^ T cell co-stimulation through different TNFRSF members produces quantitatively different cytokine phenotypes. **(A)** Primary human CD8^+^ T cells transduced with the c58/c61 TCR were stimulated for 4, 8, 16 and 24 hours with plate-immobilised pMHC and ligands to TNFRSF members at the indicated doses. The production of cytokines into the culture medium supernatant was quantified by ELISA. Surface receptors were labelled with fluorescent antibodies and quantified by flow cytometry. **(B)** Representative IFN-γ dose-responses for the four measured time points, with different colours representing the indicated doses of TNFSF ligands. **(C)** Cytokine E_max_ values normalised to E_max_ without co-stimulation at 4 hours (3 independent experiments). **(D)** Rate of change of E_max_ from (C) normalised to the rate without co-stimulation at 4 hours. E_max_ values and rates of cytokine production with and without co-stimulation were compared using multiple two-tailed t-tests. Abbreviations: * = p-value < 0.05; ** = p-value < 0.01; *** = p-value < 0.001. **(E)** Representative surface expression time courses of TNFRSF members on cells stimulated with 2000 ng/well pMHC, with or without the respective ligands.

As we previously found (11), presentation of antigen in the absence of co-stimulation induced a burst of cytokine but ultimately led to adaptation, whereby T cells stopped cytokine production so that supernatant levels remained similar after 8 hours (Fig. 2B, grey line). This is observed by the constant value of E_max_ (Fig. 2C) or by the rate of change of E_max_ approaching 0 (Fig. 2D) after 8 hours without co-stimulation.

Simultaneous engagement of TCR and these co-stimulatory receptors revealed different quantitative phenotypes (Fig. 2B). We found that 4-1BB co-stimulation had the strongest amplification on cytokine production, and this amplification continued to increase over time beyond 8 hours with the maximum cytokine production observed at the final time point (Fig. 2C). This was achieved by maintaining a high rate of cytokine production (Fig. 2D). Although CD27 co-stimulation amplified cytokine production, it appeared to be less effective at halting adaptation so that the amplification remained similar after 8 hours with the rate of change of cytokine production decreasing after this time (Fig. 2B–D). We observed similar effects on TNF and IL-2 production (Fig. S1), although IL-2 levels decreased over time, which is likely a result of consumption (discussed further below). In comparison, GITR and OX40 had almost no effect on cytokine production (Fig. 2B–D). Together, this data suggested that the effect of CD27 co-stimulation is rather front-loaded, i.e. increasing the early response within the first 8 hours only, whereas 4-1BB co-stimulation is most effective at later timepoints when T cells without co-stimulation would already halt their response.

This pattern of cytokine production was consistent with the temporal expression pattern of these receptors (Fig. 2E), with CD27 being highly expressed on resting T cells and rapidly downregulated upon engagement with its ligand CD70, while 4-1BB is not present on resting T cells and only up-regulated upon TCR-dependent activation. This activation-induced expression of 4-1BB appears to somewhat compensate for the 4-1BBL-induced downregulation of the receptor, allowing it to remain on the cell surface for longer, possibly resulting in the more persistent effect of 4-1BB co-stimulation. GITR and OX40 are similarly activation-induced, however with much slower kinetics and reaching lower levels compared to 4-1BB on CD8^+^ T cell blasts (Fig. S2), which could explain their weak effects in our system. These receptors are likely more relevant on other T cell populations, such as those among CD4^+^ T cells (32–34). Therefore, our further analysis mostly focused on CD27 and 4-1BB.

In addition to impacting the rate and duration of cytokine production, TNFRSF co-stimulation also appeared to enhance the T cell sensitivity to pMHC antigen. We observed that 4-1BBL or CD70 co-stimulation led to a 2- to 10-fold lower EC_50_ (Fig. S3). Once again, the timing of these effects matched the expression pattern of the co-stimulatory receptors: The EC_50_ initially remained unaffected by 4-1BB co-stimulation and gradually decreased over time in comparison to the control without co-stimulation (Fig. S3A,B), while the effect of CD70 was largest at the earlier timepoints and is appreciably reduced by 24 hours (Fig. S3C,D). While the effects of 4-1BB and CD27 on T cell sensitivity are statistically significant, they are dwarfed by those of adhesion receptors such as CD2 (3, 4), for which we observe up to several hundred-fold reduction in EC_50_ in our system (Fig. S3E). Therefore, we focused on understanding their impact on rate and duration of cytokine production.

### Cytokine response in adapted T cells can be rescued by co-stimulation through TNFRSF

In the previous experiments, TNFRSF co-stimulation was provided at the outset. This raises the question of whether TNFRSF co-stimulation can revert T cell adaptation once it has already been established.

While CD27 co-stimulation was not able to prevent adaptation in the time course experiments (Fig. 2), under the assumption that this is merely due to rapid downregulation of CD27, preserving CD27 expression on adapted T cells may render them receptive to CD70-induced rescue of cytokine responses. To study this, we pre-stimulated T cells with a dose-response of pMHC alone for 16 hours to induce unresponsiveness. Transferring these pre-treated cells to the same dose-range of pMHC alone for another 8 hours resulted in only modest cytokine production, confirming that these T cells had adapted to the antigen dose they experienced in the first stimulation. Engaging CD27 in this second stimulation could indeed rescue the cytokine response confirming that CD27 can rescue adapted T cells (Fig. 3A–D). In addition, the presence of CD70 in the first stimulation did not prevent unresponsiveness to pMHC alone in the second stimulation confirming that CD27 cannot override adaptation. Moreover, it rendered CD27 co-stimulation in the second phase less effective, which is expected due to the early downregulation of CD27 by CD70 in the first stimulation. Taken together, ligation of CD27 can revert T cell adaptation in a pMHC-dependent manner once it is established provided that CD27 has not been downregulated.

**Figure 3:**
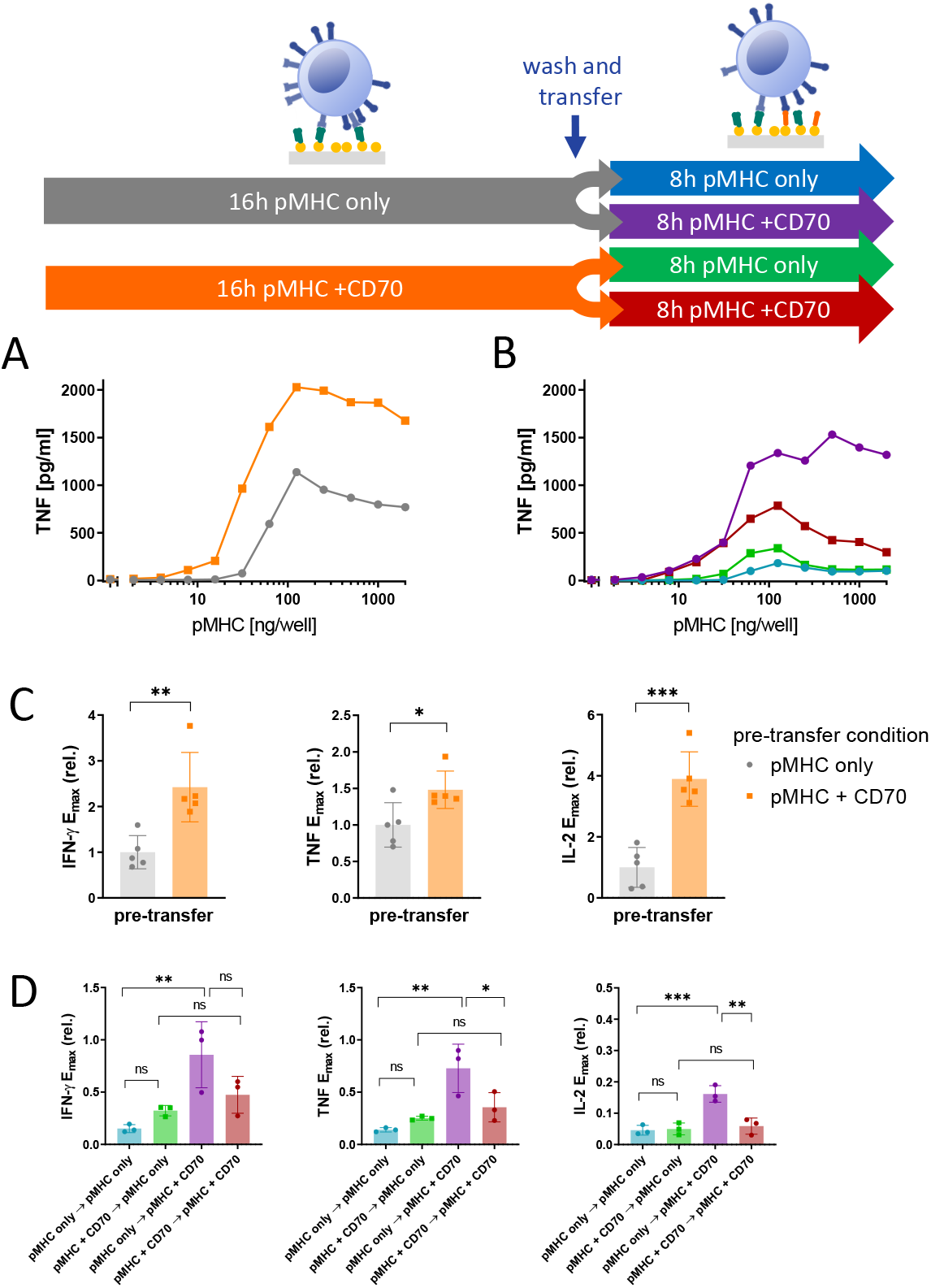
CD27 co-stimulation is capable of rescuing the cytokine response in already adapted T cells if not engaged during the first stimulation. Primary human CD8^+^ T cells transduced with the c58/c61 TCR were stimulated for 16h with pMHC doses varying from 0 to 2000 ng/well in presence or absence of 200 ng/well CD70. Cells were harvested, washed and stimulated for further 8h with identical pMHC doses which they were adapted to, with or without addition of 200 ng/well CD70. The production of the cytokines IFN-γ, IL-2 and TNF into the culture medium supernatant was quantified by ELISA. **(A)** T cell response during the first 16h stimulation from one representative experiment. **(B)** T cell response during the secondary 8h stimulation from the same experiment. In **(C)** and **(D)**, E_max_ values from three independent experiments were extracted from dose-response curve fits and normalised to the cytokine response during the 16h pre-stimulation without co-stimulation. Pre-transfer conditions (pMHC with or without CD70) in (C) were compared with a two-tailed t-test. Conditions in (D) were compared using one-way ANOVA with Šídák’s correction for multiple comparisons. Abbreviations: ns = p-value > 0.05; * = p-value < 0.05; ** = p-value < 0.01; *** = p-value < 0.001.

Although GITR co-stimulation produced only modest increases in cytokine production in the time-course, we found that in this two phase experiment it was able to partially revert unresponsiveness in pre-stimulated T cells (Fig. S4). This might have been due to the small effect of GITR being masked by the early co-stimulation-independent burst of cytokine in the time course experiments. We noted that unlike IFN-γ and TNF, GITR co-stimulation could not rescue the IL-2 response in these plate-transfer experiments and this could be explained by the ability of T cells to consume IL-2 (Fig. S4C). T cells indeed upregulated CD25 upon activation in our system (Fig. S6A), which is part of the high-affinity IL-2 receptor (35, 36).

We have recently shown that 4-1BB engagement can rescue cytokine responses from adapted T cells (11), albeit using different stimulation times. We therefore repeated the experiments using the timings above for CD27 and GITR and confirmed that delaying 4-1BB engagement can rescue cytokine responses (Fig. S5A–C). Given that 4-1BB expression is induced by pMHC-dependent TCR signalling, we could not rule out that 4-1BB expression alone was sufficient to induce cytokine production (i.e. that T cells were now licensed to secrete cytokine independent of pMHC) because in conditions without pMHC or with low doses of pMHC in the second stimulation phase, 4-1BB was not expressed. Therefore, in a second set of experiments, the T cells were all pre-stimulated with a fixed dose of pMHC and CD70 for 16 hours to uniformly induce high 4-1BB expression with minimal TCR downregulation before transferring them to a dose-range of pMHC, with or without 4-1BBL, for another 8 hours (Fig. S5D–F). The pMHC dose-dependency of the cytokine response in the second phase indicated that cytokine production was still strictly dependent on TCR stimulus, since at low pMHC doses (or at no pMHC) no response was observed despite high 4-1BB expression and 4-1BBL availability (red and orange lines), confirming 4-1BB as a bona-fide co-stimulatory receptor incapable of inducing a T cell response on its own.

The validity of conclusion drawn from these plate-transfer experiments hinges on a high standard of reproducibility of plate coatings and the maintenance of a constant pMHC stimulus. To demonstrate this, we quantified the coated amount of pMHC on the plates using an immunofluorescence assay, both before addition of cells (with or without the cognate TCR) and after a 16 hour stimulation phase and harvest of the cells for the transfer. No changes in coating were detected between the different conditions, indicating that the pMHC is not decaying or consumed at any rate relevant to the time scale of our experiments (Fig. S6B). Moreover, viability staining of the cells after the entire process of the experiment shown in Fig. S5D–F shows that the unresponsiveness of adapted T cells was not due to cell death (Fig. S6C), in addition to the fact that their response could be rescued with 4-1BB co-stimulation.

Taken together, co-stimulation by 4-1BB, CD27, and GITR can rescue cytokine production in a pMHC-dependent manner by unresponsive T cells. This is consistent with their shared signalling pathways (15) and supports the hypothesis that differences in their quantitative phenotypes is a result of surface receptor expression and regulation.

### A mechanistic model reproduces the different TNFRSF co-stimulation phenotypes based on shared signalling but different feedback control of receptor expression

We have previously published a simple mechanistic mathematical model which could explain T cell adaptation as a consequence of TCR downregulation (11). This ordinary-differential-equation (ODE) model included pMHC binding to the TCR that induced both TCR downregulation and TCR signalling (effectively an incoherent feedforward loop) that could turn on a digital switch that activated downstream signalling leading to cytokine production (Fig. 4A).

**Figure 4:**
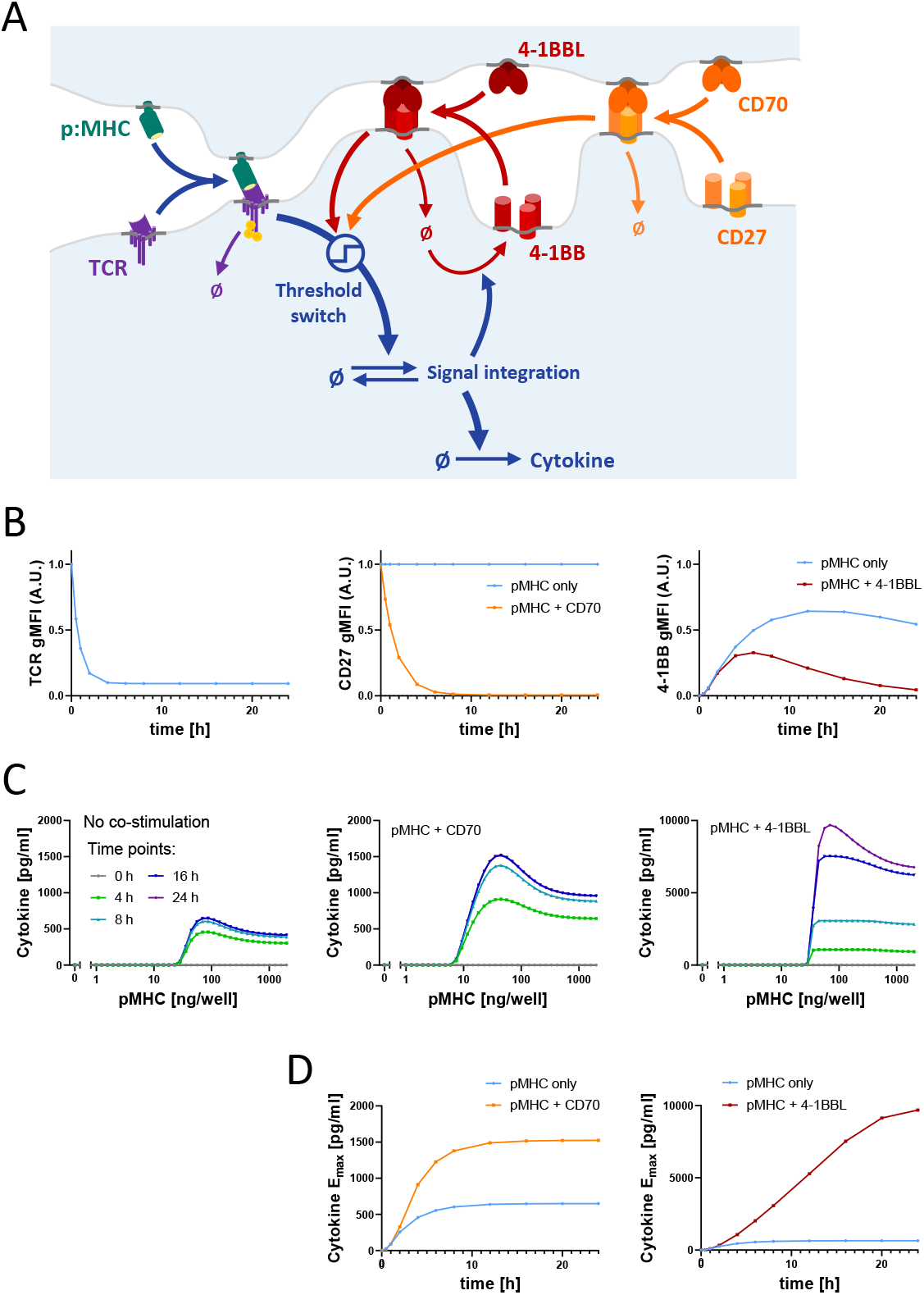
An operational mathematical model explains the divergent phenotypes of 4-1BB and CD27 based on a shared signalling mechanism but different surface receptor regulation. **(A)** Model schematic: T cell receptor (TCR) and peptide-major histocompatibility complex (pMHC) form a receptorligand complex that induces the cytokine response, gated by a threshold switch. At the same time, ligand binding causes downregulation of the TCR. Similarly, the co-stimulatory TNFRSF/TNFSF receptor-ligand pair form a complex which causes modulation of the T cell activation threshold, as well as downregulation of the TNFRSF. In the case of CD27, the receptor is present from the start, whereas 4-1BB expression is induced by TCR signalling. **(B)** Simulated time courses of TCR (left), CD27 (middle) and 4-1BB (right) surface expression using the model in (A) for pMHC in absence or presence of the respective TNFRSF ligands. **(C)** Simulated cytokine dose-responses for the indicated time points in absence (left) or presence of CD70 (middle) or 4-1BBL (right). **(D)** Simulated time courses of cytokine E_max_ in absence or presence of CD70 or 4-1BBL.

We next systematically explored where in this pathway can 4-1BB and CD27 integrate their signals to reproduce the cytokine data we had collected (Fig. S7–8). For example, we found that if CD27 and 4-1BB modulated the pMHC-TCR interaction they would only shift the dose-response EC_50_ in our model and leave the large changes in cytokine production rate and duration unexplained (Fig. S7A). We could reproduce the changes in cytokine production by allowing them to modulate TCR expression and downregulation but this required that CD27 and 4-1BB co-stimulation increase surface TCR levels. However, measurements of TCR surface expression show rather the opposite, i.e. a 4-1BBL and CD70 dose-dependent enhancement of TCR downregulation (Fig. S7B), possibly as a result of their modest abilities to increase adhesion to the stimulation surface. Using data from Fig. 3A and B, we were also able to exclude five alternative models with downstream integration points which were inconsistent with our observations (Fig. S8).

This process of elimination led us to a single model that can explain the cytokine data based on 4-1BB and CD27 signalling modulating the molecular switches that convert the analogue antigen signal into the reported digital cytokine response on single cell level (37, 38) (see Fig. 4A). Simulation of a time course using this model, with the co-stimulatory receptor being present from the start and rapidly downregulated upon engagement, was able to reproduce the CD27 co-stimulation phenotype with an increased rate of initial cytokine production but eventual arrest of the response within the same time scale of T cell adaptation in the absence of co-stimulation (Fig. 4B to D). In the case of 4-1BB, with the co-stimulatory receptor being absent on resting cells and induced upon activation, the model was also able to replicate the longer duration and hence higher rate of cytokine production at time points after 8 hours.

In summary, our mechanistic model confirmed that the difference in receptor expression are sufficient to explain the divergent phenotypes of CD27 and 4-1BB co-stimulation (Fig. 4D) when they share the same signalling mechanism. In contrast to CD27 which is short-lived due to ligand-induced downregulation, 4-1BB co-stimulation is sustained through a feedback loop which replenishes surface 4-1BB expression by activation-induced synthesis of 4-1BB.

### Sequential engagement of CD27 and 4-1BB produces synergistic effects

The operational model that we inferred from our data predicted synergy between CD27 and 4-1BB. This prediction is based on the inference that CD27 integrates its signal within the positive feedback that drives 4-1BB surface expression (Fig. 4A). Thus, the model suggests that CD27 would not only increase cytokine production directly but would also increase 4-1BB expression and therefore, improve the subsequent co-stimulation effects induced by 4-1BB engagement. We illustrated this by simulating the two-phase stimulation assay, whereby T cells were stimulated with a dose-range of pMHC with or without addition of CD70 for 16 hours, followed by a transfer to the same pMHC doses in absence or presence of 4-1BBL (Fig. 5). Not only does CD27 co-stimulation increase cytokine production in the first phase as shown previously (Fig. 2 and 3A); the model additionally predicted that 4-1BB surface expression would be elevated, and the amount of cytokine produced during the second stimulation would also be increased in the presence of 4-1BBL (Fig. 5B). These predictions were confirmed experimentally (Fig. 5C and D). Since this synergy is based on the activation-induced expression of 4-1BB, other activation-induced co-stimulatory receptors should hypothetically behave similarly. Indeed, we confirmed a similar synergy for GITR showing that IFN-γ production was significantly enhanced when T cells received early CD27 co-stimulation before GITR co-stimulation (Fig. S9).

**Figure 5:**
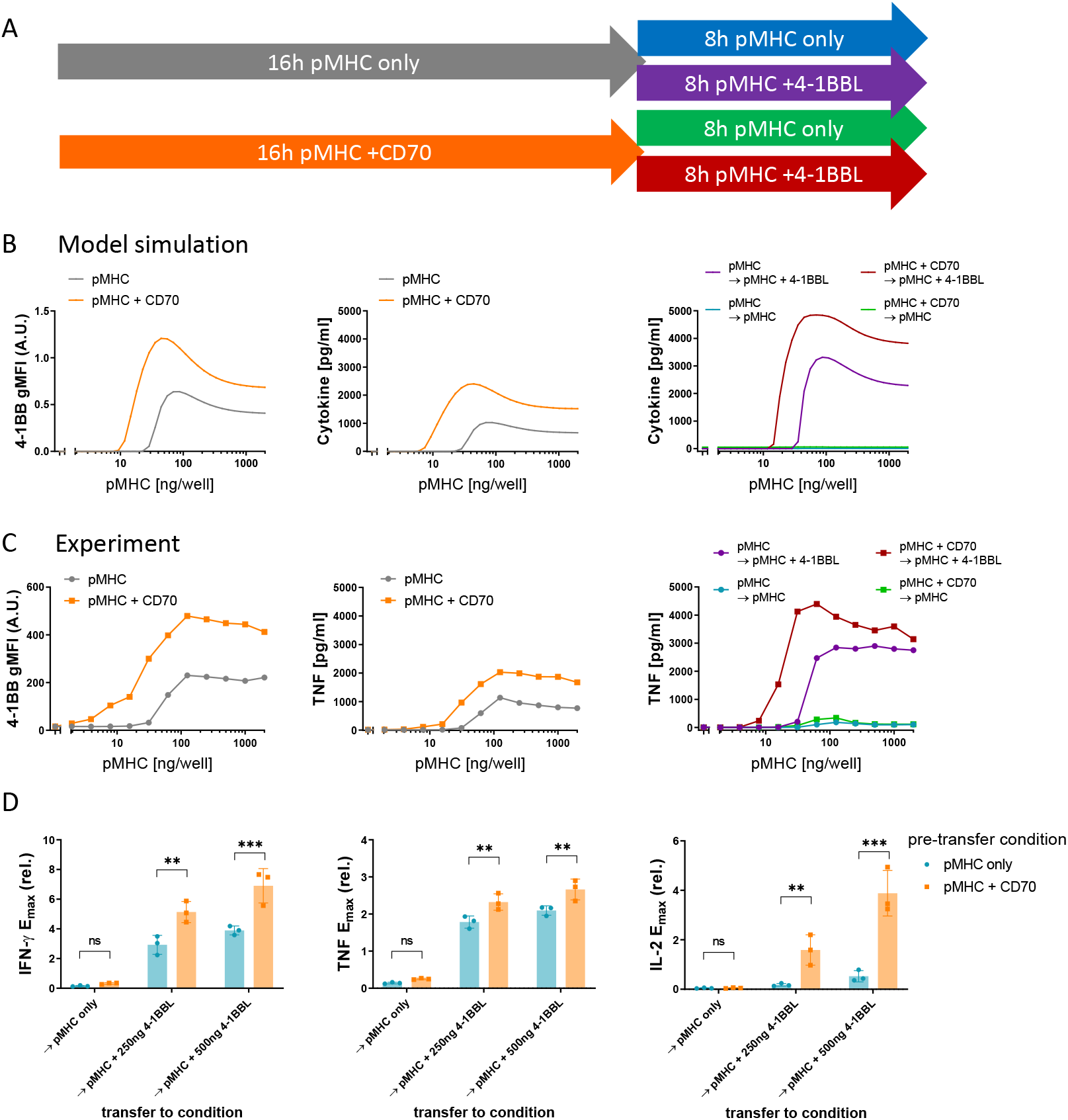
Sequential co-stimulation through CD27 and 4-1BB exhibits synergy. **(A)** Overview of experiments to explore the impact of CD27 co-stimulation on subsequent 4-1BB co-stimulation. **(B)** Predictions of the operational model (as described in Fig. 4) for the two phase stimulation experiments shown in (A). 4-1BB expression (left) and cytokine response (middle) were simulated for a 16-hour stimulation with pMHC only (blue) or presence of 200 ng/well CD70 (orange). Afterwards, the model predicted cytokine levels from these cells upon transfer to identical pMHC doses for another 8 hours, with or without addition of 500 ng/well 4-1BBL (right). The simulated experiment was performed in (C) and (D) using primary human CD8^+^ T cells transduced with the c58/c61 TCR. The production of the cytokines IFN-γ, IL-2 and TNF into the culture medium supernatant was quantified by ELISA. **(C)** T cell response during the first 16h stimulation from one representative experiment (middle) and during the secondary 8h stimulation from the same experiment (right). Cells from designated duplicate samples in the same experiment were stained with fluorescent anti-4-1BB antibodies after the first 16h stimulation and analysed by flow cytometry (left). **(D)** E_max_ values from three separate repeats of the experiment were extracted from dose-response curve fits and normalised to the cytokine response without co-stimulation during the 16h pre-stimulation. Post-transfer conditions in were compared using one-way ANOVA with Šídák’s correction for multiple comparisons. Abbreviations: ns (not significant) = p-value > 0.05; * = p-value < 0.05; ** = p-value < 0.01; *** = p-value < 0.001

## Discussion

In this study, we collected time series data with precisely titrated inputs in a minimal *in vitro* system to characterise the quantitative impact of TNFRSF co-stimulation on the T cell cytokine response. We found that while both 4-1BB and CD27 co-stimulation increased efficacy of the response with minor increases in sensitivity, only 4-1BB was able to prolong the response duration. We found only minor changes in cytokine production by GITR and OX40 co-stimulation in this initial screen. These phenotypes were consistent with their surface expression dynamics, with CD27 being rapidly downregulated upon ligand engagement, whereas 4-1BB was upregulated by TCR signalling, which partially counteracted its ligand-induced down-regulation. Both GITR and OX40 were induced with slower kinetics and to a lesser extent compared to 4-1BB. An operational model of T cell activation could explain the different phenotypes of CD27 and 4-1BB by a shared signalling mechanism (lowering the TCR signalling threshold for cytokine production) but differences in the regulation of their surface expression.

This is consistent with the current molecular view that suggests conserved signalling pathways between TRAF-binding members of the TNFRSF. While they may utilise different TRAF variants, they are commonly strong activators of NF-*κ*B and engage MAPK pathways (1), supporting a shared co-stimulation mechanism. TCR signalling is known to efficiently activate NFAT, while the other major transcription factors of T cell activation, AP-1 (through the MAPKs ERK and JNK) and NF-*κ*B, typically require stronger or prolonged stimulation (39–41). By supplementing AP-1 and NF-*κ*B activation, it is thus plausible that co-stimulation through TNFRSF can lower the amount of TCR signalling required to elicit a response. Therefore, when T cells adapt to a pMHC stimulus by downregulating their TCR signalling machinery, sufficient co-stimulation would be able to revert this state of unresponsiveness.

Interestingly, other hyporesponsive T cell phenotypes such as anergy and exhaustion are also characterised by diminished TCR signalling, either through downregulation of signalling molecules or expression of co-inhibitory receptors (42, 43). Moreover, imbalanced activation of transcription factors in favour of NFAT is also implicated in the induction of these states (40, 41, 44), so it is not surprising that TNFRSF co-stimulation has been reported to counteract both anergy and exhaustion *in vivo* (45–47). However, a causal relationship of this effect with increase in NF-*κ*B and AP-1 activation upon TNFRSF activation remains to be formally established.

While our coarse-grained ‘operational’ model captures the signal-processing behaviour of the cell, it is unable to pinpoint molecular interactions between TCR and co-stimulatory signalling. Therefore, it should be seen as complement to molecular maps rather than their replacement (24). One of the advantages of operational models is the capacity to produce predictions of T cell responses to more complex inputs, such as combinations of co-stimulatory ligands. We experimentally validated the prediction of synergy between TNFRSF members that relied on the feedback regulation of the expression of 4-1BB and other inducible TNFRSF co-stimulatory receptors. In fact, CD27 is expressed on resting T cells and is rapidly downregulated upon ligand contact, whereas expression of the other TNFRSF members has variable kinetics and requires activation of the T cell to be induced, separating their primary time of action. Feedback regulation of the inducible TNFRSF members is thus possibly capable of imprinting a ‘history’ of past activatory and co-stimulatory encounters, allowing even early co-stimulators such as CD27 to affect the later response long after they have been downregulated, if the appropriate ligands to inducible TNFRSF members are present.

Assuming that our observation is the result of feedback regulation of inducible co-stimulatory receptors expressed in proportion to the strength of T cell activation, rather than immediate integration of signals from multiple receptors, this synergy is not necessarily exclusive to the TNFRSF. It is very likely that co-signalling receptors from other protein families can affect expression of inducible TNFRSF, as well. Based on similar observations of tightly regulated transient and temporally staggered expression of co-stimulatory and co-inhibitory receptors, Chen and colleagues have proposed a ‘tidal model’, where immune cells at different stages of the response and their differentiation will be affected by different sets of co-signalling ligands (48). Our data demonstrates that these changes can be highly dynamic, allowing for rapid fine-tuning on the time scale of hours.

Adoptive cell therapies using T cells expressing chimeric antigen receptor (CAR) are now routinely used in the clinic to treat B cell malignancies. However, long-term remission is not achieved in a large fraction of patients and the therapy has yet to be proven in solid tumours (49, 50). Importantly, the ability of adoptively transferred CD8^+^ T cell blasts to produce the cytokine IFN-γ is critical for tumour control (51). Our model suggests that 4-1BB can be critical for the sustained production of IFN-γ when the TCR signalling machinery is downregulated as T cells adapt to a TCR stimulus. Therefore, inserting 4-1BB co-stimulatory domains directly into a CAR might limit its effectiveness, as CARs are also known to be downregulated upon triggering (11, 52), and we have shown that co-stimulatory receptors that are downregulated alongside the TCR (such as CD27) are unable to prevent unresponsiveness through adaptation. Targeted stimulation of endogenous 4-1BB might provide a better way to enhance the potency of T cells used for adoptive cell therapy. In a direct comparison, T cells co-transduced with a second-generation CAR and 4-1BBL for co-stimulation in *cis* and *trans* have been shown to be superior to third-generation CAR T cells *in vivo* (47). Alternatively, ‘armored’ CAR T cells engineered to inducibly produce pro-inflammatory cytokines in order to adjust the tumour microenvironment in their favour have been proposed (53). Here, type-I interferons could be promising, since they are known to induce TNFSF ligands on APCs (54).

Co-signalling receptors are critically important regulators of T cell responses and currently, these receptors are largely classified into the binary categories of co-stimulation or co-inhibition (1). Using systematic experiments, we have been able to go beyond this binary classification by identifying different quantitative phenotypes induced by co-stimulation through different TNFR members. An operational model for how T cell responses are regulated by the TCR allowed us to explore how different quantitative co-stimulation phenotypes can arise depending on where in the TCR signalling pathway a specific co-stimulation receptor integrates and how the surface expression of the co-stimulation receptor is regulated. Although these models lack molecular information, they can be helpful to provide mechanisms (how inputs are converted into outputs) and can make operational predictions (24, 55). Indeed, the model predicted how co-stimation from one receptor (CD27) can impact subsequent co-stimulation from other receptors (4-1BB, GITR). By combining quantitative experiments and mathematical modelling, it may be possible to produce an operational map for how different surface receptors quantitative control T cell responses.

## Materials & Methods

### Protein production

pMHCs were refolded *in vitro* from the extracellular residues 1–287 of the HLA-A*02:01 α-chain, β2-microglobulin, and NY-ESO-1_157-165_ peptide variant SLLAWITKV as described previously (27). TNFSF ligand expression constructs were a kind gift from Harald Wajant (Würzburg, Germany) and contained a Flag tag for the purification and a tenascin-C trimerization domain (56). We added an N-terminal AviTag as biotinylation site using standard cloning techniques. The protein was produced by transient transfection of HEK 293T cells with X-tremeGENE HP Transfection Reagent (Roche), according to the manufacturer’s instructions, and purified following a published protocol (56), with the exception of the elution step in which we used acid elution with 0.1 M glycine-HCl at pH 3.5. The pMHC or co-stimulatory ligand was then biotinylated *in vitro* by BirA enzyme, according to the manufacturer’s instructions (Avidity Biosciences), purified using size-exclusion chromatography with HBS-EP (pH 7.4, 10 mM HEPES, 150 mM NaCl, 3 mM EDTA, and 0.005% v/v Tween20) as flow buffer, and stored in aliquots at −80°C.

### Production of lentiviral particles for transduction

HEK 293T cells were seeded into six-well plates before transfection to achieve 50-80% confluency on the day of transfection. Cells were co-transfected with the respective third-generation lentiviral transfer vectors and packaging plasmids using Roche X-tremeGENE HP (0.8 µg lentiviral expression plasmid, 0.95 µg pRSV-Rev, 0.37 µg pMD2.G, 0.95 µg pMDLg/pRRE per well). The supernatant was harvested and filtered through a 0.45-µm cellulose acetate filter 24-36 h later. The affinity-matured c58c61 TCR (25) was used in a standard third-generation lentiviral vector with the human EF1α promoter.

### T cell isolation and culture

Up to 50 ml peripheral blood was collected by a trained phlebotomist from healthy volunteer donors after informed consent had been taken. This project has been approved by the Medical Sciences Interdivisional Research Ethics Committee of the University of Oxford (R51997/RE001), and all samples were anonymized in compliance with the Data Protection Act. Alternatively, leukocyte cones were purchased from National Health Services Blood and Transplant service. Only HLA-A2-peripheral blood or leukocyte cones were used because of the cross-reactivity of the high-affinity receptors used in this project, which leads to fratricide of HLA-A2+ T cells (57). CD8^+^ T cells were isolated directly from blood using the CD8^+^ T Cell Enrichment Cocktail (STEMCELL Technologies) and density gradient centrifugation according to manufacturer’s instructions. The isolated CD8^+^ T cells were washed and resuspended at a concentration of 1 × 106 cells/ml in culture medium (RPMI 1640 with 10% fetal bovine serum, 100 units penicillin and 100 μg streptomycin per ml) supplemented with 50 U/ml IL-2 and 1 × 10^6^ CD3/CD28-coated Human T-Activator Dynabeads/ml (Life Technologies). The next day, 1 × 10^6^ T cells were transduced with the 2.5 ml virus-containing supernatant from one well of HEK 293T cells, supplemented with 50 U/ml of IL-2. The medium was replaced with fresh culture medium containing 50 U/ml IL-2 every 2-3 days. CD3/CD28-coated beads were removed on day 5 after lentiviral transduction, and the cells were used for experiments on days 10-14. TCR expression was assessed by staining with NY-ESO 9V PE-conjugated tetramer (inhouse produced using refolded HLA*A02:01 with NY-ESO-1_157-165_ 9V and streptavidin-PE [Bio-Rad AbD Serotec or BioLegend]) using flow cytometry.

### T cell stimulation

T cells were stimulated with titrations of plate-immobilized pMHC ligands with or without co-immobilized ligands for accessory receptors. Ligands were diluted to the working concentrations in sterile PBS. 50 μl serially diluted pMHC were added to each well of high-binding capacity streptavidin-coated 96-well plates (#15500; Thermo Fisher Scientific). After a minimum 45-min incubation at room temperature, the plates were washed with sterile PBS. Where accessory receptor ligands were used, those were similarly diluted and added to the plate for a second incubation. After washing the stimulation plate with PBS, 7.5 × 10^4^ T cells were added in 200 μl culture medium (RPMI 1640 with 10% fetal bovine serum, 100 units penicillin and 100 μg streptomycin per ml) without IL-2 to each stimulation condition. The plates were spun at 50–100 ×g for 1 min to settle down the cells and then incubated at 37°C with 5% CO_2_. At the indicated time points of the stimulation experiments, cells were harvested by pipetting and transferred to V-bottom 96-well plates. The harvested cells were pelleted (5 min at 520 ×g) and processed for flow cytometry, and the supernatants were collected for ELISAs. To stimulate T cells in two phases with different conditions, stimulation plates for a second condition were prepared as described above. At the time point of transfer between conditions, cells were harvested, pelleted in V-bottom 96-well plates, and subsequently resuspended in fresh pre-warmed culture medium (200 μl/well) before transferring them to the prepared stimulation plates for the second phase. To settle down the cells again, plates were briefly spun at 50–100 ×g for 1 min and returned to the incubator (37°C / 5% CO_2_) for the specified times.

### Flow cytometry

Flow cytometry was used to quantify surface expression of TCR, co-stimulatory receptors and activation markers at specified time points and at the end of stimulation experiments. Harvested cells were pelleted in V-bottom 96-well plates (5 min at 520 ×g) and kept on ice during the staining procedure. The pellets were resuspended in staining buffer (PBS with 1% BSA) containing fluorescently labelled pMHC tetramers and/or fluorophore-conjugated antibodies (Table) at previously titrated working concentrations (usually 1:200 for commercially available antibodies from BioLegend). After 20 min, cells were washed twice with staining buffer (200 well, 5 min at 520 ×g) and resuspended in 80 μl/well PBS for flow cytometry. Samples were analysed on a BD FACSCalibur™ or BD LSRFortessa™ X-20 with a BD High Throughput Sampler for automated acquisition. Flow cytometry data was analysed in FlowJo V10.

### ELISA

After harvesting the cells from the stimulation experiments, the supernatants were separated from the cell pellets, collected in round-bottom 96-well plates and kept on ice for short-term storage (< 12 hours). To quantify the cytokines IFN-γ, IL-2 and TNF, ELISAs were performed using Nunc MaxiSorp™ flat-bottom 96-well plates and the respective Invitrogen™ Uncoated ELISA Kits (ThermoFisher Scientific) according to the manufacturer’s protocol. A BioTek ELx405 plate washer was used for washing steps, and absorbance at 450 nm and 570 nm was measured using a SpectraMax M5 plate reader (Molecular Devices). Standards in duplicates were included for each plate to generate calibration curves for the calculation of cytokine concentrations.

### Data analysis

We fit the following bell-shaped function to each dose-response curve using the function *lsqcurvefit* in Matlab (Mathworks, MA),

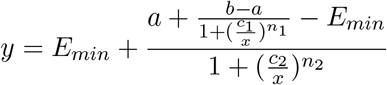

We then used the smooth line produced by this function with 10,000 points to directly calculate the maximum amount of cytokine produced across different pMHC concentrations (E_max_) and the pMHC concentration producing the half-maximal response (EC_50_) for each dose-response curve (Supplementary Fig. S1A). We note that these are not fitted parameters but rather metrics determiend directly from the line produced by the bell-shaped function. In certain cases, the cytokine response was so weak that the technical noise in the ELISA made it unfeasible to accurately estimate the EC_50_. Therefore, EC_50_ values were excluded for dose-responses where the fit E_max_ was below 50 pg/ml, which was usually the case for IL-2 at late time points. To further minimise the contribution of technical variability between experiments (originating, for instance, from differences in quality of stimulatory ligands, transduction efficiency of the cells and donor variability) the extracted absolute E_max_ and EC_50_ values were normalised to the mean E_max_ and EC_50_ values of each readout across all conditions in each experiment. Therefore, pooled and averaged E_max_ and EC_50_ values were not plotted as absolute values but as relative fold-change compared to one standard condition (e.g. pMHC without co-stimulation), as indicated for each figure. Statistical tests with appropriate corrections for multiple testing were performed as indicated for each figure in GraphPad Prism 8.

### Mathematical modelling

We have used a mathematical model of pMHC binding to the TCR that induces both TCR downregulation and digital cytokine activation. The model presented in this work is a simplified version of a model we have recently described (11). The model is represented by a system of ordinary-differential-equations (ODEs). Reaction rates in the model are described by mass-action kinetics with the exception of receptor-ligand binding, which is simplified under the assumption that equilibrium is reached rapidly (seconds to minutes) compared to the timescale of the experiments (hours), and the digital behaviour of TCR signalling described by an error function erf to account for natural variability of the activation threshold within the cell population. The model was numerically integrated using the solver *ode23s* in Matlab (Mathworks, MA). The following set of ordinary differential equations describes the base model:

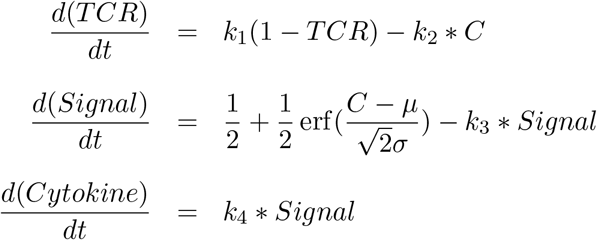

Here, the parameter *μ* is the mean activation threshold in a population of T cells, *σ* is its standard deviation of *μ* in the population, and *C* is the amount of pMHC-TCR complexes,

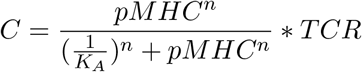

where *K*_*A*_ is the affinity and *n* is the hill number. The surface expression of co-stimulatory receptors was modelled as follows:

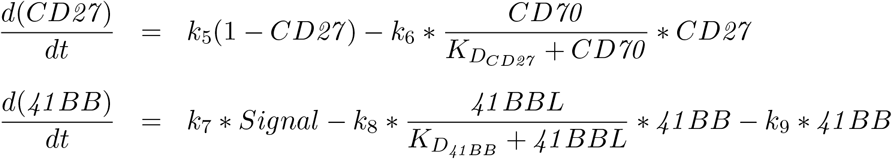

Under the assumption that co-stimulation through CD27 and 4-1BB affect the T cell response by modulating the activation threshold, the equation for the integrated signal changes as follows:

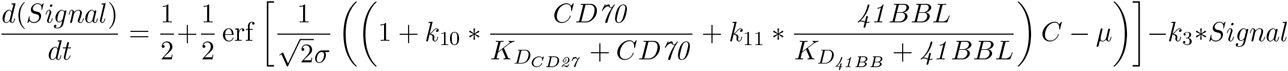

The parameter values for the generation of Figs. 4 and 5 using this assumption are listed in Table S1.

The 6 alternative models for the CD27 co-stimulation transfer experiments in Fig. S8 were simulated with the following equations, respectively, that were altered from the base model (above) to integrate co-stimulation at a different position:

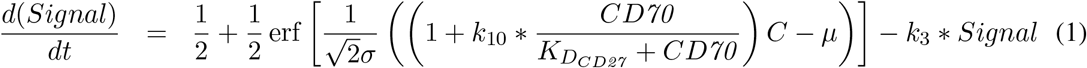

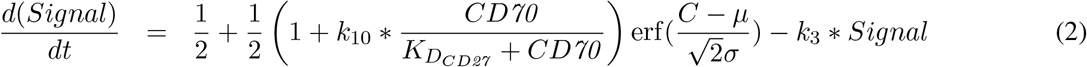

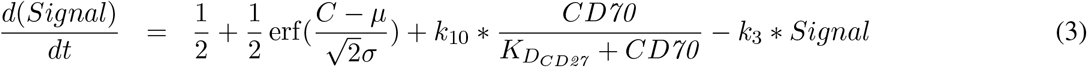

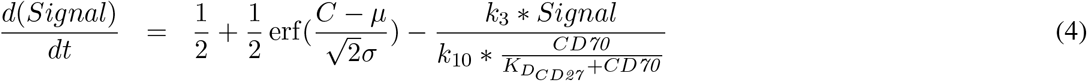

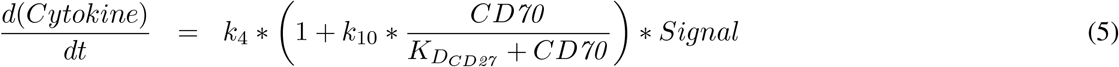

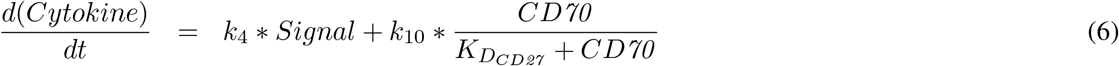

## Acknowledgement

We thank Harald Wajant for providing expression plasmids for all TNFRSF ligands, Simon J. Davis for providing expression plasmid for CD58, and Adaptimmune Ltd for providing the c58c61 TCR. We thanks P. Anton van der Merwe and Marion H. Brown for helpful discussions.

## Supplementary Figures

**Figure S1:**
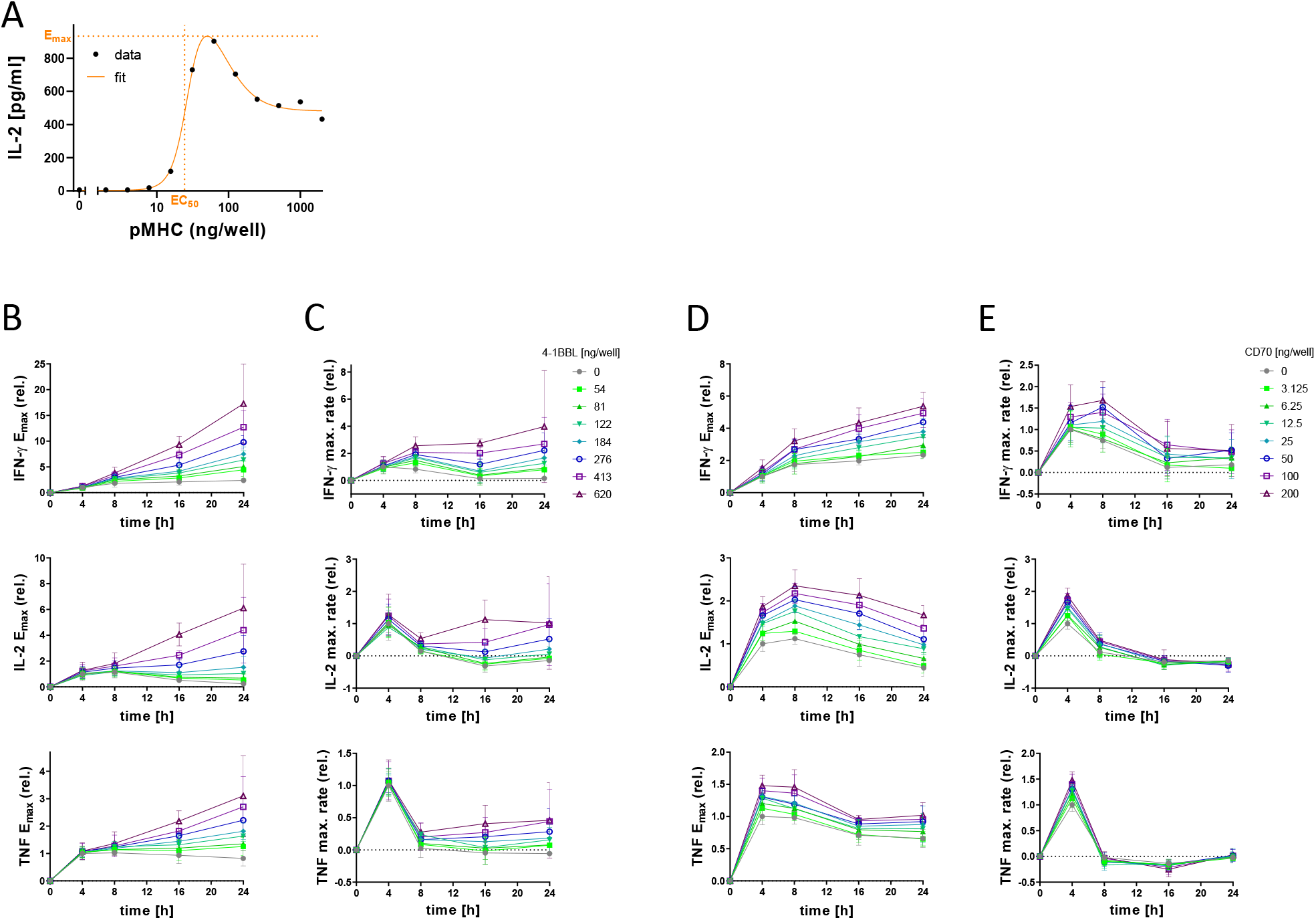
4-1BB co-stimulation delays adaptation of the T cell cytokine response, whereas CD27 co-stimulation only increases early cytokine production. Primary human CD8^+^ T cells transduced with the c58/c61 TCR were stimulated for 4, 8, 16 and 24 hours with plate-immobilised pMHC, with or without 4-1BBL in (B) and (C) or CD70 in (D) and (E) at the indicated doses. The production of the cytokines IFN-γ, IL-2 and TNF into the culture medium supernatant was quantified by ELISA. **(A)** Cytokine dose-response curve fit and extraction of metrics. As an example, an IL-2 dose-response data set was fit with a bell-shaped function as described in the Methods. Graphically shown as dotted lines are the E_max_ and EC_50_ which were empirically determined. In **(B)** and **(D)**, cytokine E_max_ values from three independent experiments were extracted from dose-response curve fits and normalised to the amount of cytokine produced without co-stimulation at the 4 h time point. In **(C)** and **(E)**, average rates of cytokine production were calculated from the extracted E_max_ values and normalised to the rate of cytokine production in the first 4 hours without co-stimulation.

**Figure S2:**
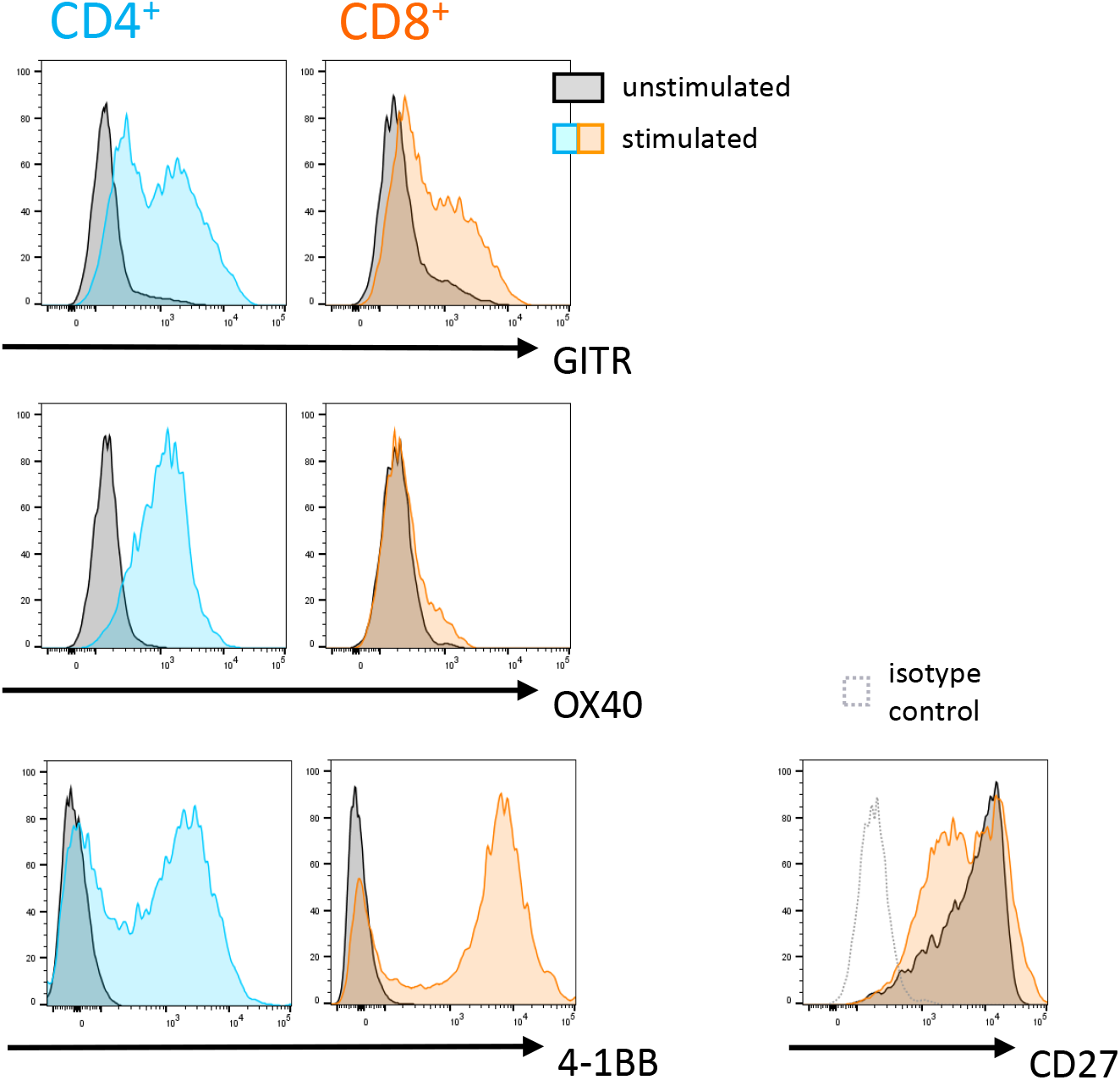
GITR and OX40 expression is induced on CD8^+^ T cells to a lesser extent than on CD4^+^ T cells. Expanded primary human CD4^+^ and CD8^+^ T cell blasts (day 10-13) were stimulated with plate-immobilised anti-CD3 antibodies (clone UCHT1, 1μg/ml) for 18 hours. The indicated surface receptors were labelled with fluorescent antibodies and analysed by flow cytometry. Shown here are flow cytometry histograms for unstimulated (grey) and stimulated (blue/orange) T cells.

**Figure S3:**
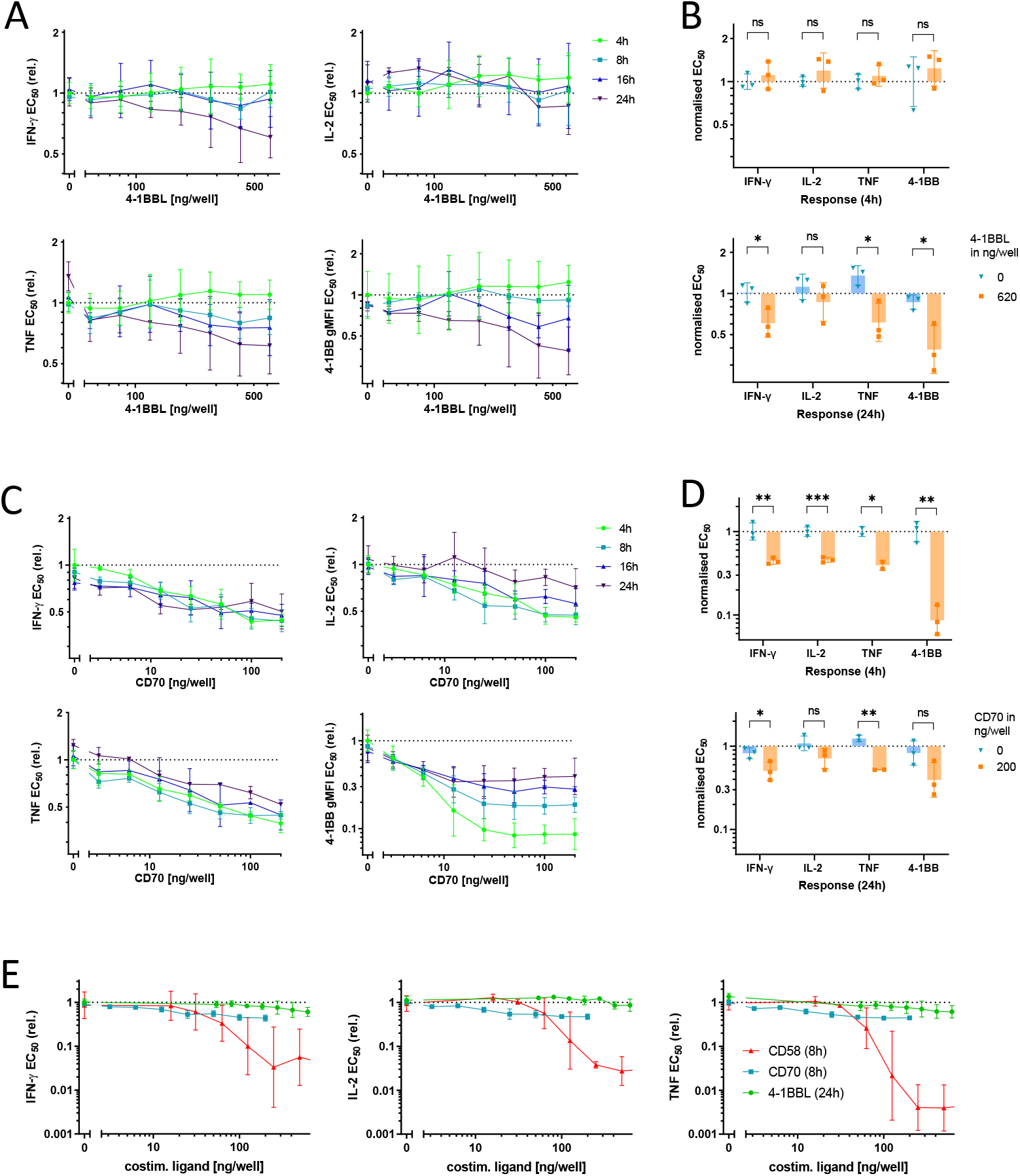
Co-stimulation through TNFRSF members lowers the EC_50_ of the T cell response to pMHC. Primary human CD8^+^ T cells transduced with the c58/c61 TCR were stimulated for 4, 8, 16 and 24 hours with plate-immobilised pMHC, with or without 4-1BBL in (A) and (B) or CD70 in (C) and (D) at the indicated doses. The production of the cytokines IFN-γ, IL-2 and TNF into the culture medium supernatant was quantified by ELISA. Surface 4-1BB was labelled with a fluorescent anti-4-1BB antibody and quantified by flow cytometry. In **(A)** and **(C)**, EC_50_ values from three separate experimental repeats were extracted from dose-response curve fits and normalised to the 4-hour time point without co-stimulation. To quantify statistically significant effects on sensitivity, EC_50_ values at 4 or 24 hours were compared in **(B)** and **(D)** with two-tailed t-tests between the conditions with or without co-stimulation. For comparison, the same experiment was carried out using CD58 as co-stimulatory ligand to CD2 **(E)**. EC_50_ values from three separate experimental repeats were extracted from dose-response curve fits, normalised to the 4-hour time point without co-stimulation and plotted along with selected data from (A) and (C). Abbreviations: ns (not significant) = p-value > 0.05; * = p-value < 0.05

**Figure S4:**
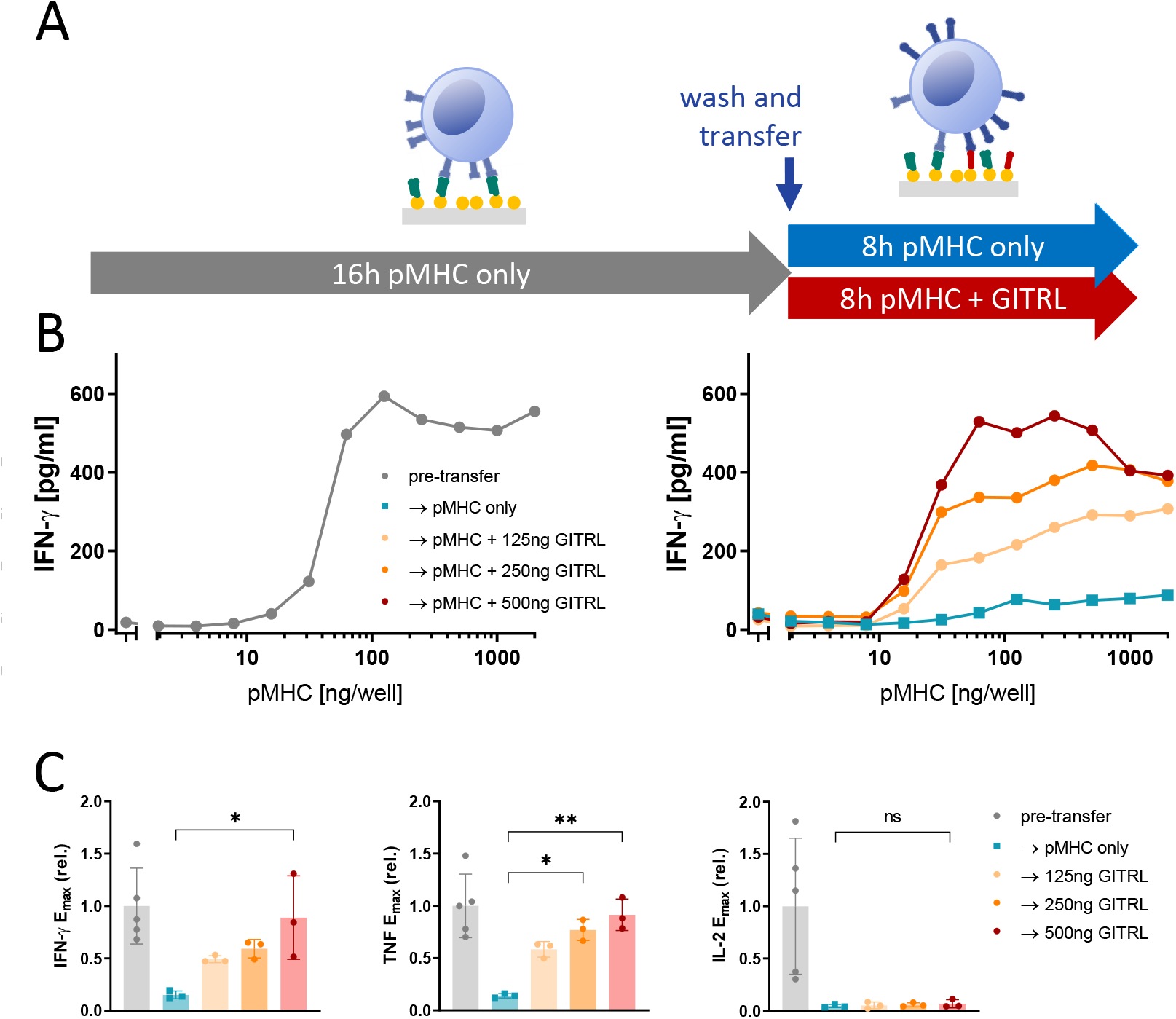
GITR co-stimulation is capable of partially rescuing the cytokine response in already adapted T cells. **(A)** Schematic of the experimental design for (B) and (C). Primary human CD8^+^ T cells transduced with the c58/c61 TCR were stimulated for 16h with pMHC doses varying from 0 to 2000 ng/well. Cells were harvested, washed and stimulated for further 8h with identical pMHC doses which they were adapted to, in absence or presence of GITRL. The production of the cytokines IFN-γ, IL-2 and TNF into the culture medium supernatant was quantified by ELISA. **(B)** T cell response during the first 16h stimulation from one representative experiment (left) and during the secondary 8h stimulation from the same experiment (right). **(C)** E_max_ values from three separate experimental repeats were extracted from dose-response curve fits and normalised to the cytokine response during the 16h pre-stimulation. Post-transfer conditions were compared using one-way ANOVA with Šídák’s correction for multiple comparisons. Abbreviations: ns = p-value > 0.05; * = p-value < 0.05; ** = p-value < 0.01

**Figure S5:**
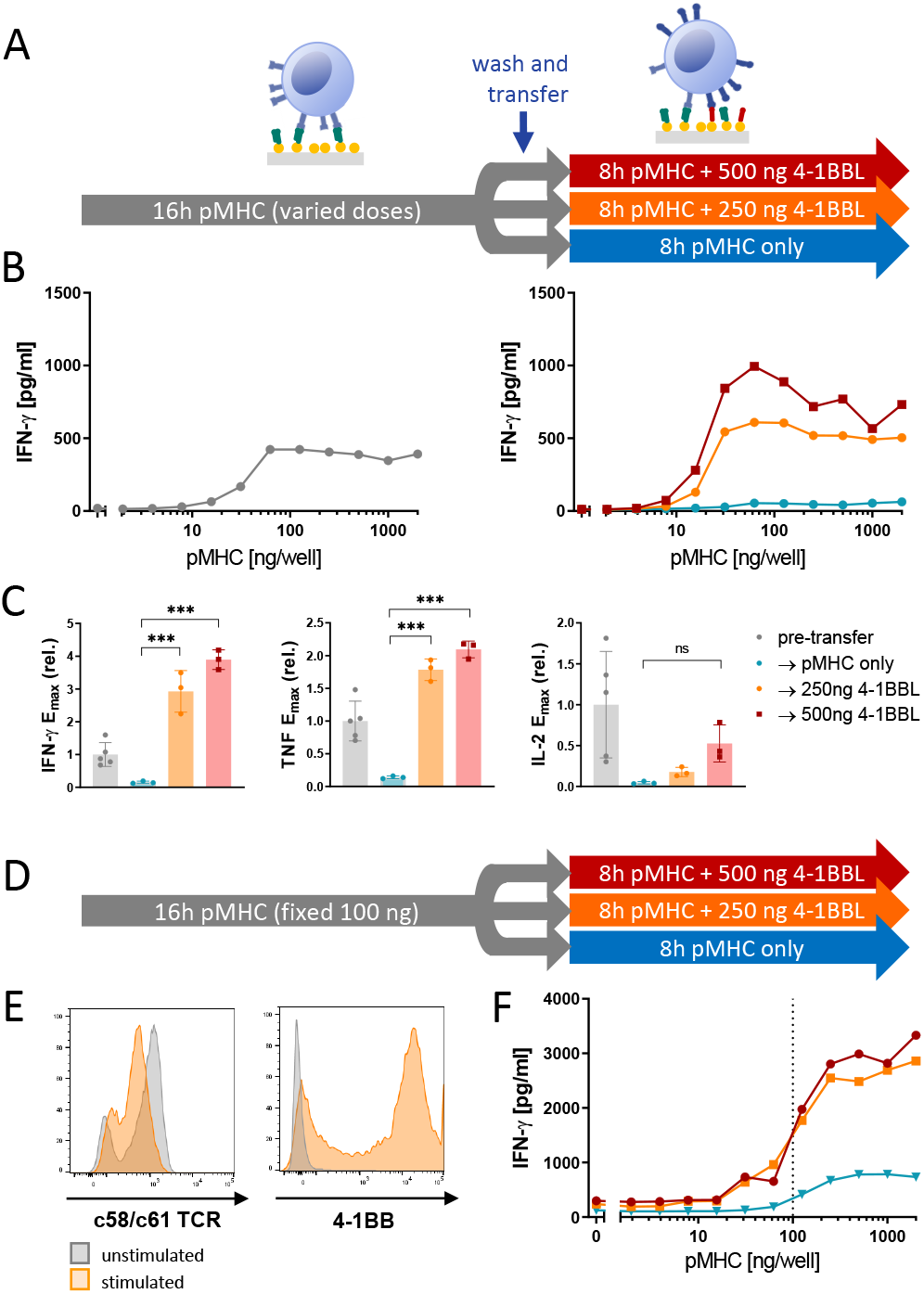
4-1BB co-stimulation rescues the cytokine response in already adapted T cells in a TCR-signalling-dependent manner. **(A)** Schematic of the experimental design. Primary human CD8^+^ T cells transduced with the c58/c61 TCR were stimulated for 16h with pMHC doses varying from 0 to 2000 ng/well. Cells were harvested, washed and stimulated for further 8h with identical pMHC doses which they were adapted to, in absence or presence of 4-1BBL. The production of the cytokines IFN-γ, IL-2 and TNF into the culture medium supernatant was quantified by ELISA. **(B)** T cell response during the first 16h stimulation from one representative experiment (left) and during the secondary 8h stimulation from the same experiment (right). **(C)** E_max_ values from three separate experimental repeats were extracted from dose-response curve fits and normalised to the cytokine response during the 16h pre-stimulation. Post-transfer conditions were compared using one-way ANOVA with Šídák’s correction for multiple comparisons. **(D)** For (E) and (F), T cells were uniformly stimulated for 16h with pMHC and CD70 (each at 100 ng/well) in order to induce 4-1BB expression with minimal TCR downregulation. Cells were harvested, washed, and stimulated for a further 8h with pMHC and 4-1BBL at the indicated doses. **(E)** TCR and 4-1BB expression on cells before and after the 16h pre-stimulation from one representative experiment out of four. **(F)** T cell response during the secondary 8h stimulation from one representative experiment out of four. The production of the cytokines IFN-γ, IL-2 and TNF into the culture medium supernatant was quantified by ELISA. Abbreviations: ns = p-value > 0.05; * = p-value < 0.05; ** = p-value < 0.01; *** = p-value < 0.001.

**Figure S6:**
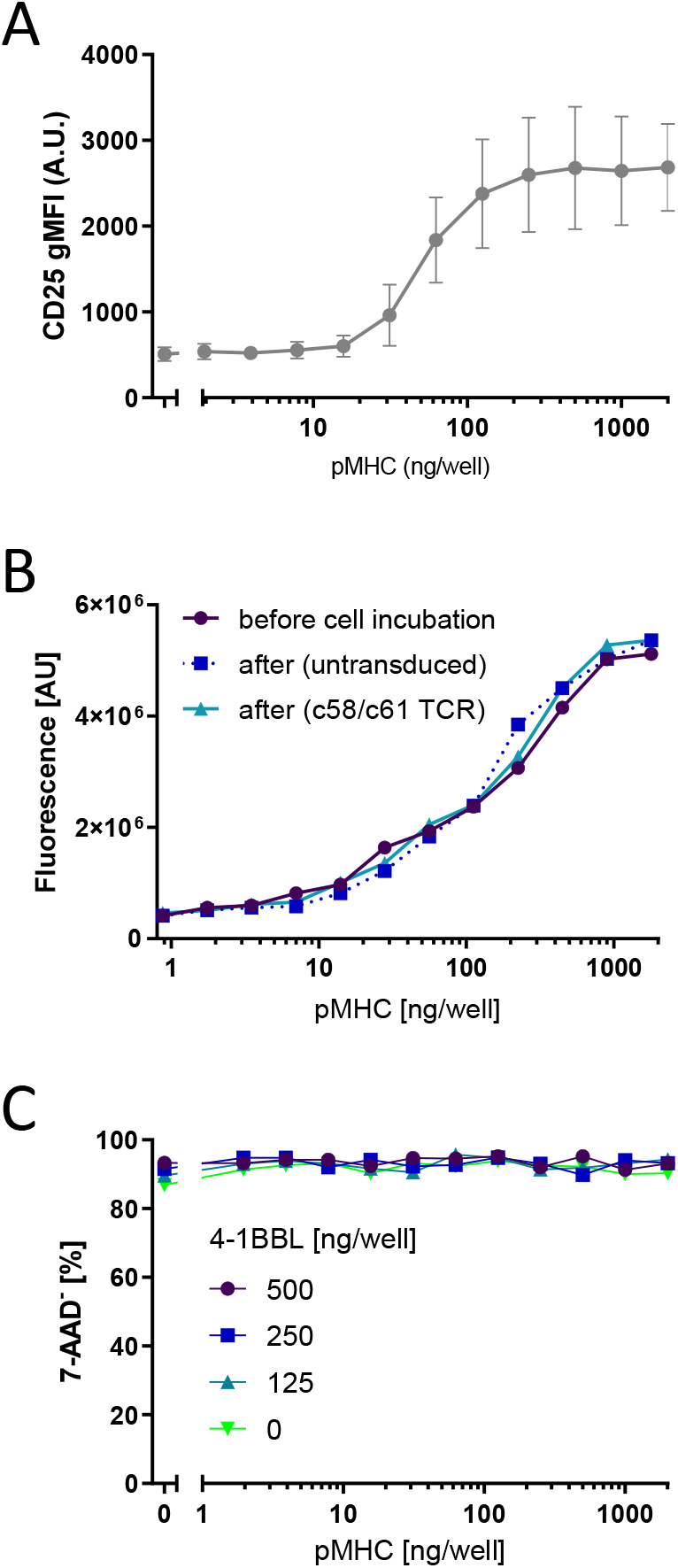
Testing limitations of the plate-based experimental system. **(A)** T cell activation induces the expression of CD25, the IL-2 receptor α-chain. Primary human CD8^+^ T cells transduced with the c58/c61 TCR were stimulated for 16 hours with pMHC at the indicated doses. Surface CD25 was labelled with a fluorescent anti-CD25 antibody and quantified by flow cytometry. Shown here is the averaged doseresponse curve for 3 experimental repeats. **(B)** Harvesting cells from coated plates after stimulation does not dislodge detectable amounts of plate-bound pMHC. Densities of pMHC immobilised on streptavidin-coated plates were quantified using a conformation-sensitive anti-HLA-A/B/C antibody (clone w6/32) either before or after incubation with primary human CD8^+^ T cells (either transduced with the c58/c61 TCR or untransduced) for 16h. **(C)** Transfer of cells between stimulation plates does not cause notable levels of cell death. Primary human CD8^+^ T cells transduced with the c58/c61 TCR were stimulated for 16h with pMHC and CD70 (each at 100 ng/well). Cells were harvested, washed, and stimulated for further 8h with pMHC and 4-1BBL at the indicated doses. Cell death after the experiment was quantified by flow cytometry using 7-AAD staining.

**Figure S7:**
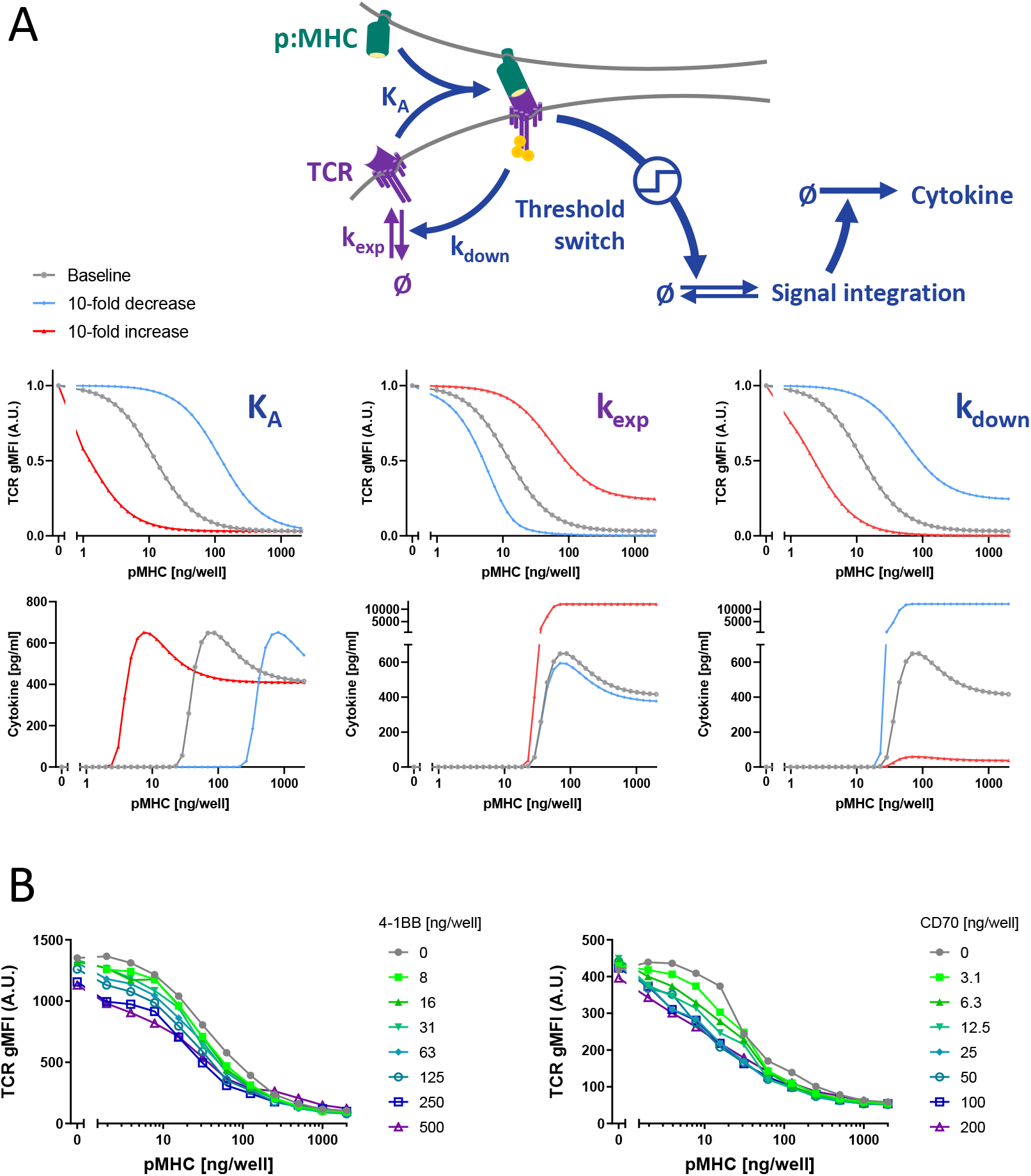
TNFRSF co-stimulation phenotypes cannot be simulated by modulation of TCR expression, downregulation or avidity to pMHC. As in the base model of Figure 4, T cell receptor (TCR) and peptide-major histocompatibility complex (pMHC) form a receptor-ligand complex that induces the cytokine response, gated by a threshold switch. At the same time, ligand binding causes downregulation of the TCR. **(A)** Potential effects of modulation of TCR-pMHC association constant (*K*_*A*_), TCR expression rate (*k*_*exp*_) and the rate of induced TCR downregulation (*k*_*down*_) by co-stimulation on TCR surface expression and cytokine production were simulated as 10-fold increases or decreases of the respective parameters. **(B)** Primary human CD8^+^ T cells transduced with the c58/c61 TCR were stimulated for 8 hours with plate-immobilised pMHC and ligands to TNFRSF members at the indicated doses. Surface TCR was labelled with fluorescent pMHC tetramers and quantified by flow cytometry.

**Figure S8:**
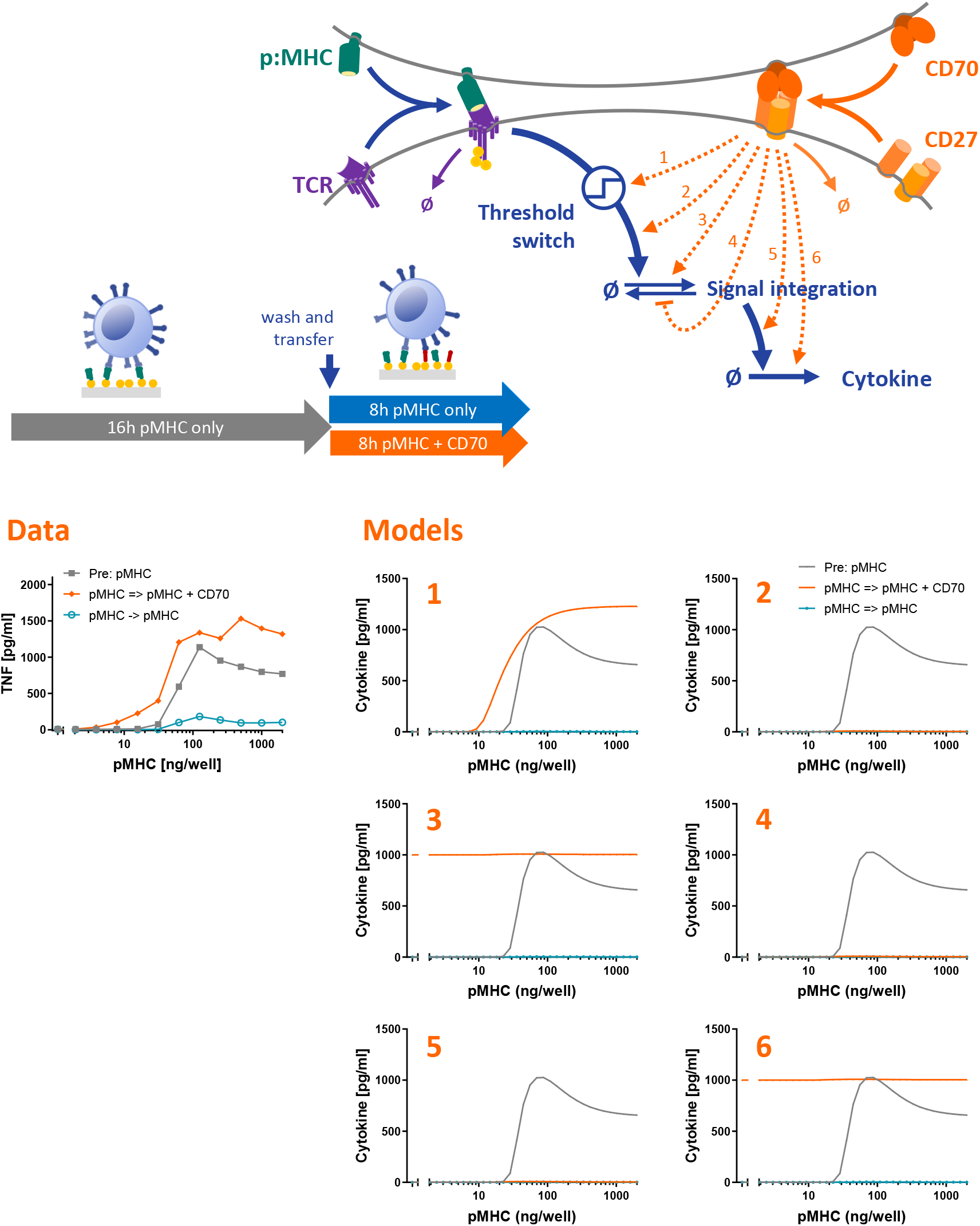
The effect of TNFRSF co-stimulation on the T cell response in transfer experiments can be explained by modulation of the activation threshold. The model in Fig. 4 is used to simulate the transfer experiment (see Data) for 6 different integration points for CD27 co-stimulation. As observed in the edxperimental data, a pMHC-dependent T cell response in the second stimulation (orange curves) is only observed in model 1 when CD27 lowers the activation threshold of the switch.

**Figure S9:**
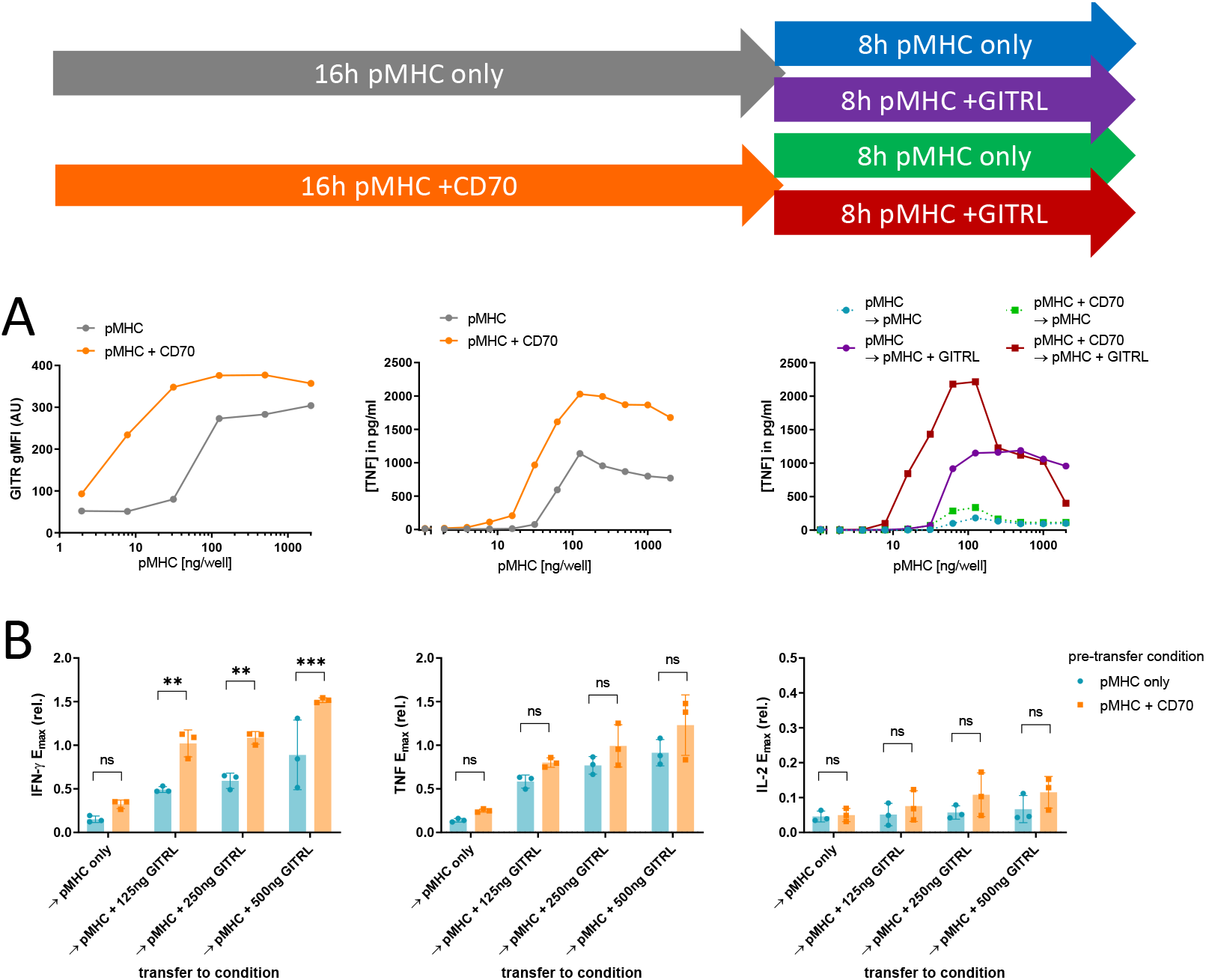
Co-stimulation through CD27 and GITR synergise. Primary human CD8^+^ T cells transduced with the c58/c61 TCR were stimulated for 16h with pMHC doses varying from 0 to 2000 ng/well in presence or absence of 200 ng/well CD70. Cells were harvested, washed and stimulated for further 8h with identical pMHC doses which they were adapted to, with or without addition of 500 ng/well GITRL. The production of the cytokines IFN-γ, IL-2 and TNF into the culture medium supernatant was quantified by ELISA. **(A)** T cell response during the first 16h stimulation from one representative experiment (middle) and during the secondary 8h stimulation from the same experiment (right). Cells from designated duplicate samples in the same experiment were stained with fluorescent anti-GITR antibodies after the first 16h stimulation and analysed by flow cytometry (left). **(B)** E_max_ values from three separate repeats of the experiment were extracted from dose-response curve fits and normalised to the cytokine response without co-stimulation during the 16h pre-stimulation. Post-transfer conditions in were compared using one-way ANOVA with Šídák’s correction for multiple comparisons. Abbreviations: ns (not significant) = p-value > 0.05; * = p-value < 0.05; ** = p-value < 0.01; *** = p-value < 0.001

